# Language remains strongly left-lateralized in older adults: A cross-sectional and longitudinal study

**DOI:** 10.64898/2026.07.17.739182

**Authors:** Elizabeth H.T. Chang, Anna Seydell-Greenwald, Trevor KM Day, Daniyal Nisbet, Peter E Turkeltaub

## Abstract

Language is typically supported by the left hemisphere, but the prevalence and stability of language lateralization in older adults remain unclear. Although acquired language disorders disproportionately affect older adults, most studies of language lateralization have focused on younger populations. We examined language lateralization and its relationship with age in a large cohort of healthy older adults.

115 healthy adults (57 F, 58 M; mean age 59.6 years) completed functional MRI during an adaptive semantic decision task. Lateralization indices were calculated using a bootstrap-based laterality index (LI) approach for whole-hemisphere, frontal, and temporal regions. Relationships between age and lateralization were examined using Pearsons and Bayesian correlation analyses. Fifteen participants completed repeat imaging after a mean interval of 44.2 months.

Language activation was strongly left-lateralized, with 96% of participants classified as left-lateralized and 3% as right-lateralized. Frontal and temporal LIs were strongly correlated. No significant relationship was observed between age and lateralization, and no significant longitudinal changes in lateralization were observed. Bayesian analyses supported absence of both age-related effects and longitudinal change in lateralization.

Language lateralization remains strongly left-lateralized and unchanging in healthy aging. Findings suggest that aging should not be a factor in the incidence and severity of aphasia from lateralized pathology.

## Introduction

Language is widely considered to be a left-lateralized function of the human brain. Early observations by Dax and Broca demonstrated that left hemisphere damage often results in aphasia, establishing one of the earliest links between cognition and hemispheric specialization^1,2^. This left dominance for language was later supported by pharmacological hemispheric inactivation using the intracarotid amobarbital procedure (IAP or Wada test), in which temporary disruption of the left hemisphere produces language impairment whereas right hemispheric inactivation does not^3^. Together, lesion studies and Wada testing established that language is predominately supported by the left hemisphere in most individuals.

Despite this population-level observation, some individuals appear to show atypical language lateralization. Evidence for atypical organization comes from rare cases of crossed aphasia following right hemisphere injury^4^ as well as a report of preserved language after extensive left hemisphere damage^5^. Determining prevalence of atypical language lateralization informs expectations for language outcomes after right- and left-hemisphere injury and provides a baseline for examining hemispheric shifts in language processing after injury^6^.

Overall, functional MRI (fMRI) studies in healthy adults have found that approximately 95% of right-handers are left hemisphere dominant for language^7–9^. Importantly, while most neurological disorders affecting language occur in older adults, lateralization studies have focused on younger adults, with a mean age of about 25 years old. This limitation is particularly relevant because several influential models of cognitive aging propose that hemispheric specialization may decrease with age, resulting in more bilateral organization in older adults^10^. These age-related reductions in hemispheric asymmetry are thought to reflect compensatory recruitment or reduced neural specialization^11^.

However, findings regarding age-related changes in language lateralization have been inconsistent. While some studies report less lateralized language activity emerging from young adulthood to later adulthood^9,12^, others observe age-related reductions in lateralization only within specific ROIs and subgroups such as right-handed men^13^, and others finding no relationship between age and language lateralization^14^. Furthermore, some studies reporting age-related changes in laterality undersampled older adults, the population most relevant to neurological disorders affecting language^9,12,13^. Moreover, age-related differences in task performance and cognitive effort may influence observed activation patterns, with bilateral recruitment in older adults reflecting compensatory responses to increased processing demands rather than reorganization of language networks^15^.

In the present study, we examined language lateralization in a large sample of healthy older adults using a highly reliable and strongly lateralizing semantic decision fMRI task that adaptively adjusts difficulty in order to mitigate potential confounds related to age-related differences in task performance and cognitive effort^16^. We report rates of typical and atypical lateralization and assess whether language lateralization varies as a function of age. A subset of participants also completed longitudinal follow-up imaging, allowing assessment of the stability of language lateralization over several years.

## Methods

### Participants

115 healthy adults participated in this study (mean age = 59.6 years, range = 22-84; 57 females, 58 males, mean years of education = 16.9, range 10-21), enrolled as control groups for studies of left hemisphere stroke (ClinicalTrials.gov IDs: NCT04991519, NCT06700005). Participants were native English speakers with at least 10 years of education and adequate vision and hearing. Exclusion criteria included history of significant brain injury, neurological, psychiatric, or learning disorders. Participants included 98 right-handed, 12 left-handed, and 5 ambidextrous individuals, determined using a cutoff of +/- 40 on the Edinburgh Handedness Inventory (EHI)^17^. The racial composition of the sample was 63% White, 33% Black or African American, 3% more than one race and 1% Asian. 3% of participants identified as Hispanic or Latino. A subset of 15 participants, 9 women and 6 men, (13 left-lateralized, 2 right-lateralized) completed repeat imaging sessions separated by a mean interval of 44.2 months (SD = 14.36, range 25.8-79.7 months) with a mean age of 61.4, allowing assessment of the longitudinal stability of language lateralization. Written informed consent was obtained according to the Declaration of Helsinki and study protocol was approved by the Georgetown University Institutional Review Board.

### fMRI Task

Language activation was measured using a validated adaptive semantic decision task shown to reliably activate a set of left-lateralized language regions^16^. During the language condition, participants viewed pairs of words and indicated whether they were semantically related (e.g., fish – pond) via button press. During a control condition, participants judged whether pairs of pseudofont strings were identical. Task difficulty was adaptively adjusted using a staircase procedure based on performance. Functional MRI data were preprocessed using standard procedures in AFNI^18^, and subject level activation maps were generated using a general linear model contrasting real words versus pseudofonts, and normalized to MNI space. Additional details regarding image acquisition and preprocessing can be found in Anderson et al^19^.

### Lateralization Index Toolbox and Regions of Interest

Lateralization indices were computed using an LI-toolbox (download: https://www.medizin.uni-tuebingen.de/de/das-klinikum/einrichtungen/kliniken/kinderklinik/forschung/forschung-iii/software with SPM12)^20^, implemented in SPM12 and MATLAB 2023b. The toolbox quantifies hemispheric asymmetry based on the relative distribution of task-related activation between left and right hemispheres, yielding values ranging from −1 (fully right-lateralized) to +1 (fully left-lateralized). A bootstrap-based method was used to estimate lateralization across a range of statistical threshold, reducing sensitivity to arbitrary threshold selection and the influence of outlier voxels^20^. LI estimates were generated using the toolbox’s bootstrap procedure, which resamples activation values across multiple thresholds and combines estimates using weighted averaging. All analyses were performed using default parameters unless otherwise specified. Additional details about the toolbox can be found in prior publications^20^.

Regions of interest (ROIs) were defined using meta-analytic activation maps from Neurosynth for the term “language” (https://neurosynth.org/analyses/terms/language/), thresholded at z = 3. To avoid any bias toward one hemisphere or the other, symmetric ROIs were created by taking the union of the thresholded Neurosynth map and its left-right flipped counterpart. To reduce potential confounds from medial structures, voxels within 10 +/- mm of the midline were excluded from the analysis. For regional analyses, the resulting bilateral language mask was further subdivided into frontal and temporal components by manually separating regions along the Sylvain fissure (Figure 1A). In addition to the regional analyses, whole-hemisphere LIs were calculated using all the voxels above threshold across the left and right cerebral hemispheres, excluding 10 +/- mm of the midline. Mean LIs were reported for frontal, temporal ROIs and whole-hemisphere.

**Figure 1.**
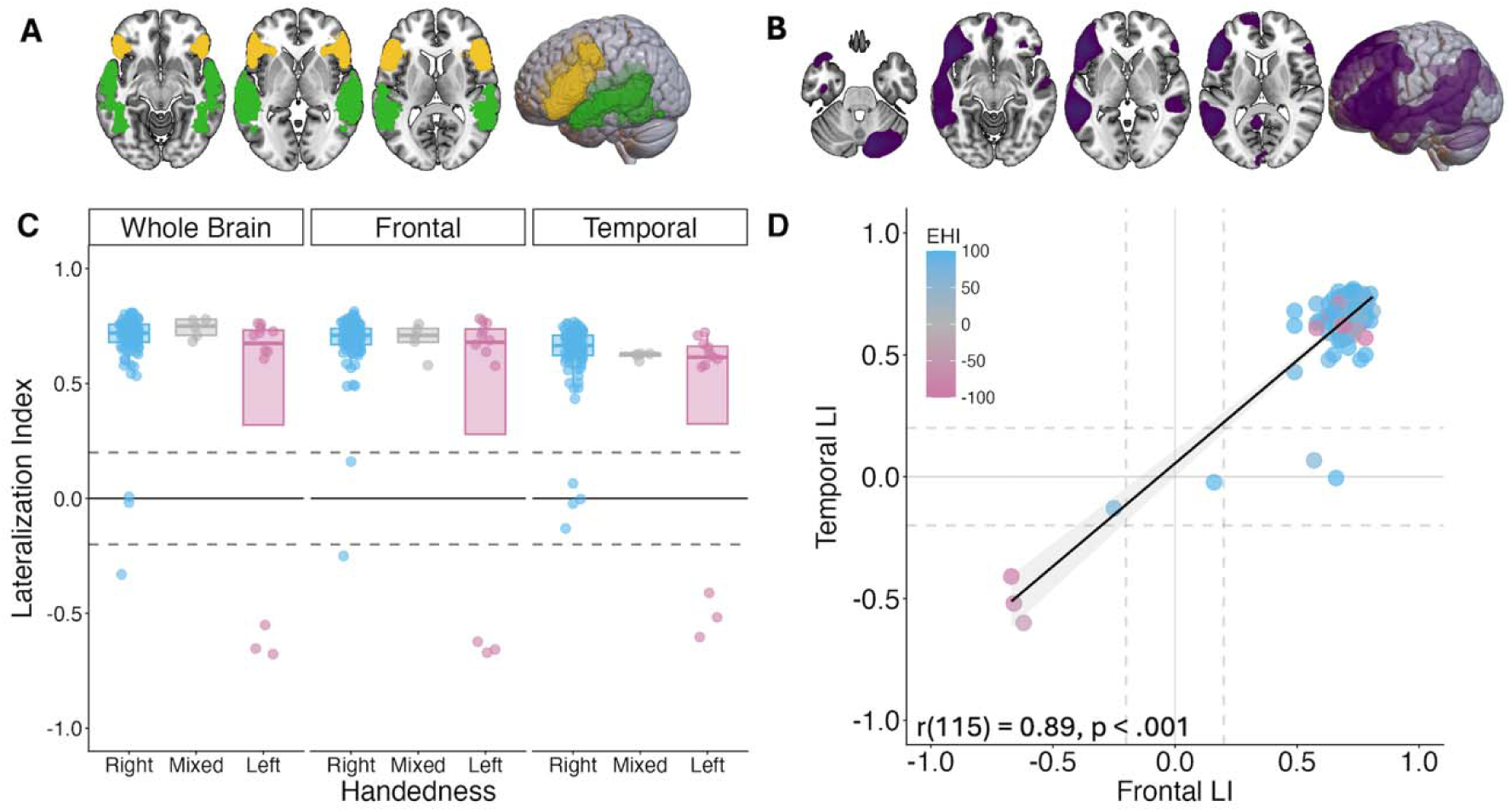
Language lateralization in healthy older adults. **(A)** Regions of interest. Frontal (yellow) and temporal (green) ROIs derived from Neurosynth language masks. Whole-hemisphere LIs (not shown) were calculated using all suprathreshold voxels within the cerebral hemispheres (excluding voxels within ±10 mm of the midline). **(B)** Penetrance map for semantic decision task. Penetrance map (purple) is thresholded at t = 3.5 and show voxels above the threshold active in >20% of participants. **(C)** Language lateralization by handedness group. Boxplots and individual participant data are shown for each ROI for right-handed, left-handed and ambidextrous participants. **(D)** Correlation between frontal and temporal LIs. Horizontal dashed lines indicate thresholds used to classify participants as left-lateralized (LI > 0.20), bilateral (−0.20 ≤ LI ≤ 0.20), or right-lateralized (LI < −0.20). Hemispheric lateralization is shown to be consistent across language regions. Each point represents an individual participant and is colored according to Edinburgh Handedness Inventory (EHI) score. Pearson correlation coefficients and associated p-values are shown.

### Statistical Analysis

Lateralization indices (LIs) were computed for each participant within the whole-hemisphere mask as well as within frontal and temporal ROIs. LIs were treated both as continuous measures and categorical variables. For categorical classification, participants were classified as left-lateralized (LI > 0.2), right-lateralized (LI < −0.2), or bilateral (−0.2 < LI < 0.2), consistent with commonly used thresholds in the literature ^21,22^.

The proportion of participants classified as left-, bilateral- and right-lateralized were calculated across whole-hemisphere, frontal and temporal analyses. Given the relationship between handedness and language lateralization^9,23,24^, these proportions were also examined in right-handed participants separately.

Pearson correlations were used to examine relationships between age and LI measures across whole-hemisphere, frontal, and temporal analyses. To further quantify evidence supporting the absence of age-related relationships, Bayesian correlation analyses were additionally performed, with Bayes factors (BF_01_) representing evidence favoring the null hypothesis of no association between age and LI. Correlation analyses were conducted across the full sample, in right-handed participants only, and right-handed participants classified as left lateralized for language. Additionally, since Nenert et al.^13^ found that age-related reductions in LI may be sex-related, linear regression models including Age, Sex and an Age × Sex interaction were fit among right-handed participants to assess whether the relationship between age and LI differed between men and women. Lastly, to assess the longitudinal stability of language lateralization, paired t-tests compared LI values between two timepoints in a subset of participants who completed repeat imaging.

## Results

Across all participants, language activation showed a strongly left-lateralized pattern (96% left-lateralized, 3% right-lateralized, 1% neither) (Figure 1B). Mean LI values were positive across all regions, including whole-hemisphere (mean = 0.70, SD = 0.11), frontal (mean = 0.65, SD = 0.24), and temporal ROIs (mean = 0.61, SD = 0.23) (Table 1). Among right-handed participants (n=98), 97% showed left-lateralized language at the whole-hemisphere level, whereas only 1% were classified as right-lateralized and 2% as bilateral (Table 1, Figure 1C). Across all participants, frontal and temporal LIs were significantly positively correlated (r=0.892, p<.001) (Figure 1D).

**Table 1.** Mean lateralization indices and categorical lateralization by handedness. Mean language lateralization indices (LI±SD) and proportions of participants classified as left-lateralized (LI > 0.20), bilateral (−0.20 ≤ LI ≤ 0.20), or right-lateralized (LI < −0.20) for frontal and temporal regions of interest and whole-hemisphere by handedness group. Positive LI values indicate greater left-hemisphere lateralization.

| ROI | Mean LI (SD) | Right-Handed (n=98) | Ambidextrous (n=5) | Left-Handed (n=12) |
| --- | --- | --- | --- | --- |
| Frontal | 0.65 (0.24) | Left: 96, Right: 1, Bilateral: 1 | Left: 5, Right: 0, Bilateral: 0 | Left: 9, Right: 3, Bilateral: 0 |
| Temporal | 0.61 (0.23) | Left: 94, Right: 0, Bilateral: 4 | Left: 5, Right: 0, Bilateral: 0 | Left: 9, Right: 3, Bilateral: 0 |
| Whole Brain | 0.70 (0.11) | Left: 95, Right: 1, Bilateral: 2 | Left: 5, Right: 0, Bilateral: 0 | Left: 9, Right: 3, Bilateral: 0 |

No significant associations were observed between age and whole-hemisphere LI across the full sample, with a Bayes factor favoring the null hypothesis of no relationship (r = - 0.15, p = 0.11, BF_01_ = 2.46) (Figure 2A). Similar results were observed when analyses were restricted to right-handed participants (r = −0.17, p = 0.10, BF_01_ = 2.09). These correlations were influenced by the few outlier participants with atypical lateralization. When analyses were further restricted to only left-lateralized participants, correlations between age and whole-hemisphere LI approached zero (r = −0.04, p = 0.57, BF_01_ = 7.60). Similar findings were observed for frontal and temporal ROIs (Table 2A).

**Figure 2.**
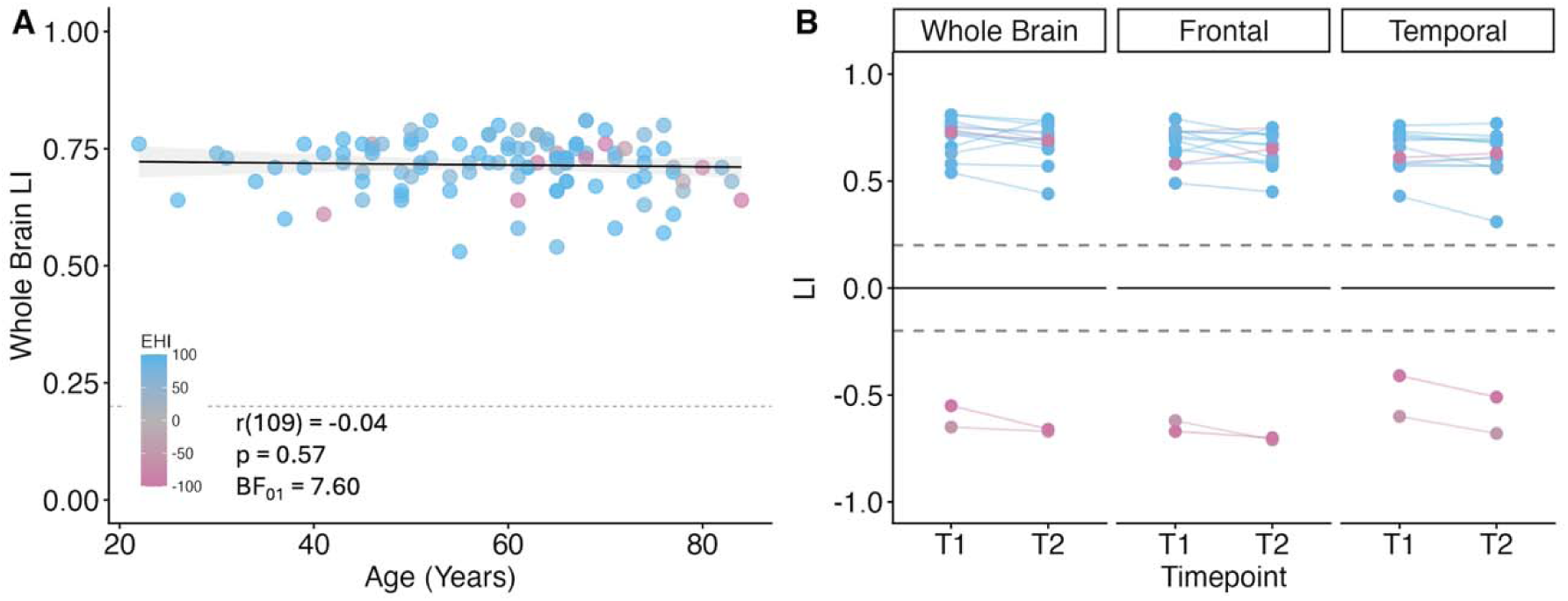
Age-related stability of language lateralization. **(A)** Relationship between age and whole-hemisphere lateralization index (LI) across left lateralized participants. No significant association was observed between age and lateralization index. Pearson correlation coefficients, p-values, and Bayes factors supporting the null hypothesis (BF_01_) are displayed on the plot. **(B)** Longitudinal stability of language lateralization. LI for whole-hemisphere, frontal and temporal ROIs shown at Timepoint 1 and Timepoint 2 for participants who completed repeat imaging. Lines connect measurements from the same participant across timepoints, illustrating within-subject changes in language lateralization over time.

**Table 2.**
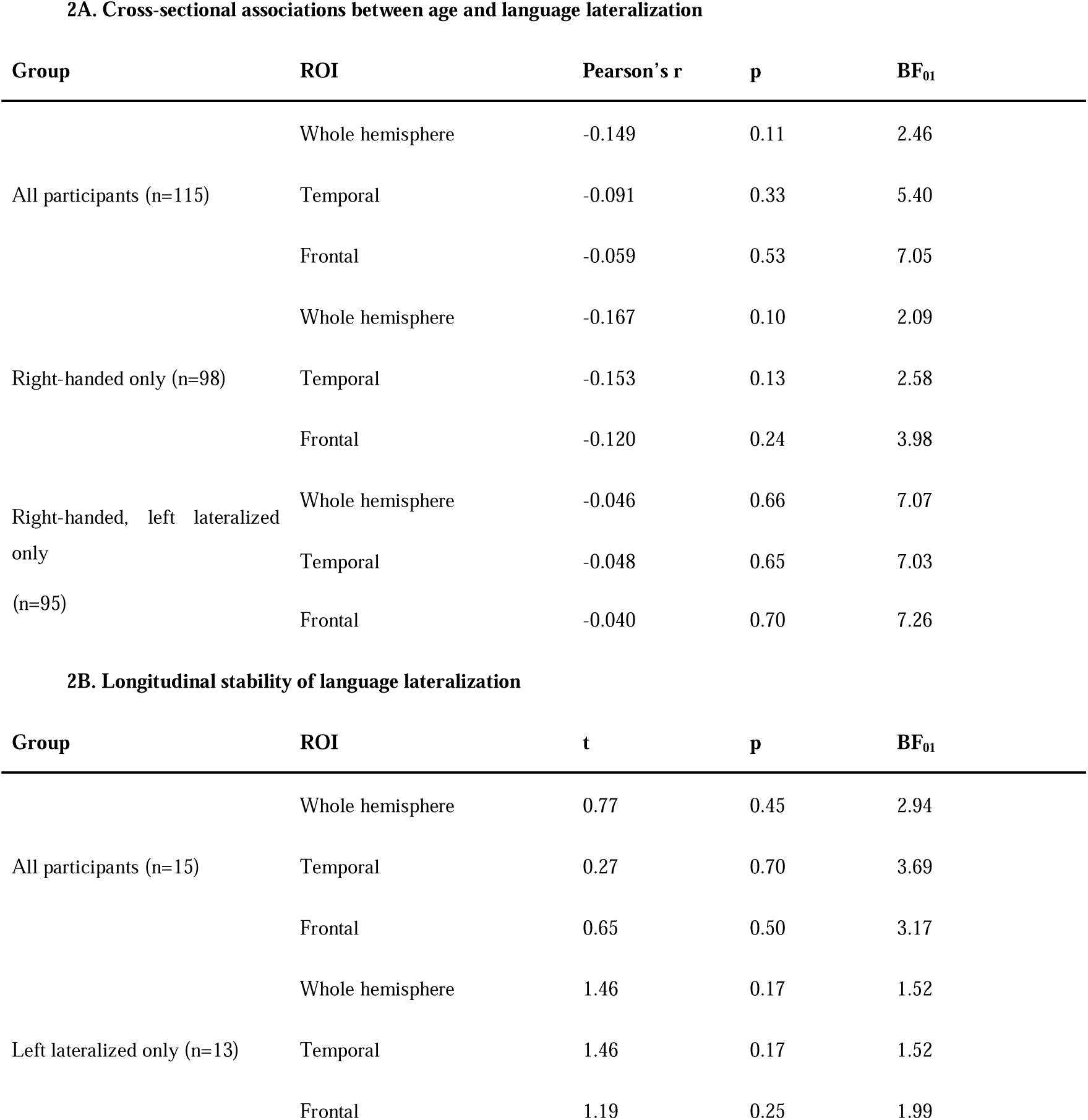
Cross-sectional and longitudinal analyses of language lateralization. (A) Cross-sectional associations between age and language LIs across whole-hemisphere, temporal and frontal regions. Bayesian correlation analyses are reported as Bayes factors favoring the null hypothesis. (B) Longitudinal stability of language lateralization in participants who completed repeat imaging. Paired t-tests compared absolute values of LIs between timepoints, with Bayes factor values indicating evidence favoring the null hypothesis of no change over time.

To assess whether age-related changes in language lateralization differed by sex, linear regression models including Age, Sex, and an Age × Sex interaction term were fit among right-handed participants. No significant Age × Sex interaction was observed for whole-hemisphere (t = −0.75, p = .457), frontal (t = 0.22, p = 0.83), or temporal LIs (t = −1.11, p = 0.27).

The longitudinal stability of language lateralization was tested in the 15 participants scanned twice two or more years apart (Figure 2B). Paired t-tests of the absolute values of the LIs in all 15 participants revealed no significant changes in LI between Timepoint 1 and Timepoint 2 for any ROI, with moderate evidence favoring the null hypothesis of no change (frontal (t(14) = 0.647, p = .50, BF_01_ = 3.17), temporal (t(14) = 0.272, p = 0.70, BF_01_ = 3.69), whole-hemisphere (t(14) = 0.77, p = 0.45, BF_01_ = 2.94)). The results remained similar when constrained to the 13 left-lateralized participants (Table 2B).

## Discussion

We examined language lateralization in a large sample of healthy older adults using a validated semantic decision fMRI paradigm and a standardized laterality index approach. Consistent with prior work in younger samples, language was strongly left-lateralized at the group level, with 97% of right-handed participants classified as left-lateralized. Frontal and temporal LI measures were highly correlated, indicating hemispheric dominance was largely consistent between language regions. Both cross sectional and longitudinal analyses yielded evidence that language lateralization does not change as a function of age within older adulthood.

The prevalence estimates of left lateralization in healthy right-handed adults observed here are slightly higher than most prior studies using Wada testing, fTCD, and fMRI on younger adults. Using a verb generation paradigm and fTCD, Knecht et al.^25^ reported left-hemisphere language dominance in 92.5% of right-handed participants. Similarly, previous fMRI studies have generally reported left-lateralization rates of approximately 92-95% in healthy adults^7–9,12,13^, but direct comparison across studies is complicated by methodological heterogeneity. For example, Pujol et al.^8^ used a silent phonemic word-generation task, whereas Szaflarski et al.^12^ employed verb generation in response to auditorily presented nouns. These tasks differ in the extent to which they engage semantic, lexical retrieval, phonological and executive processes, and consequently may vary in their ability to produce strongly lateralized activation. The nature of the control task used in fMRI contrasts may also contribute to the degree of lateralization of activity. Indeed, our mean LI of 0.70 is higher than the mean LI reported in many of these studies. Studies also employed different LI classification criteria, ranging from a directional cutoff at LI = 0^13^ to bilateral ranges of 0.1^7^, 0.2^9,12^ and 0.25^8^, further complicating direct comparison of prevalence estimates. Regardless of these differences, our findings are consistent with a conclusion that the prevalence of left lateralization of language is not lower in older adults compared with younger adults.

Contrary to predictions from models proposing age-related reductions in hemispheric specialization^10^, Bayesian analyses of both our cross sectional and longitudinal data favored the conclusion that language lateralization does not change with aging in older adults. While some previous studies have reported decreased laterality from young adulthood to late middle-age, age effects have often been inconsistent across language paradigms, brain regions, and participant subgroups. For example, Nenert et al.^13^ observed age-related reductions in laterality only within temporo-parietal regions and only among right-handed men. Springer et al.^9^ observed an inverse relationship between LI and age, but had only a few participants over age 30. Overall, our findings, supported by both cross sectional and longitudinal data, demonstrate that the most older adults have left lateralized language and that degree of laterality is unrelated to age between the ages of 30 and 85.

One possible explanation for discrepant findings in the aging literature is that apparent reductions in language lateralization may be influenced by differences in task demands rather than changes in language organization. Several studies have suggested that age-related decreases in lateralization may reflect differences in cognitive effort, sensory processing or task difficulty, due to increasing compensatory engagement of bilateral cognitive networks to accomplish tasks^15,26–28^. The task used in the present study was specifically designed to robustly engage the language network while ensuring that effort for task performance is matched across a range of ability levels through an adaptive staircase procedure^16^. This approach minimizes the potential for effort-related confounds associated with age to affect LI values.

These considerations are important because estimates of atypical language lateralization are used to interpret clinical outcomes following brain injury and reported numbers are highly variable. For example, Pedersen et al.^29^ reported 9% of a large cohort to be aphasic after right-hemisphere stroke, slightly higher than would be predicted by our findings. A more recent study found that 48.1% right hemisphere stroke survivors presented with acute aphasia as measured by the NIH Stroke Scale Best Language score, and suggested that right hemisphere language is more common than is reported in previous work^30^. The present findings instead suggest that aphasia due to right hemisphere stroke should be quite rare, and that such a high rate of aphasia after right hemisphere stroke likely reflects false positives on the NIH Stroke Scale or another source of noise in the clinical data. This discrepancy deserves further investigation using a more specific clinical measure of aphasia.

Some limitations should be acknowledged. Although the sample was large for an fMRI study of language lateralization in older adults, we under-sampled young adults under 30 years of age, limiting our ability to assess changes in lateralization in this age range. Additionally, longitudinal follow up was available in only a subset of participants and spanned about four years. Longer term follow up will be needed to determine whether language lateralization remains stable within individuals across decades of aging.

In summary, language was strong left lateralized in this large sample of healthy older adults, with atypical language organization remaining rare, particularly among right-handed individuals. Cross-sectional and longitudinal analyses provided evidence that language lateralization does not change across later adulthood. These findings suggest that hemispheric specialization for language is remarkably stable in healthy aging and provide important normative benchmark for interpreting atypical language organization in clinical populations.

## Data availability

The data in this study have recently been made publicly available through the GRAND repository. De-identified participant data can be accessed at [https://osf.io/8aqfp] and additional information about the data acquisition and preprocessing can be found in the accompanying data release paper^19^.

## Acknowledgements

We thank the following individuals who contributed to data collection, in alphabetical order: Elizabeth Dvorak, Sara Dyslin, Trini Kelly, Alycia Laks, Devna Mathur, Sachi Paul and Candace van der Stelt.

